# Enhanced Anticoagulation Activity of Functional RNA Origami

**DOI:** 10.1101/2020.09.29.319590

**Authors:** Abhichart Krissanaprasit, Carson M. Key, Kristen Froehlich, Sahil Pontula, Emily Mihalko, Daniel M. Dupont, Ebbe S. Andersen, Jørgen Kjems, Ashley C. Brown, Thomas H. LaBean

## Abstract

Anticoagulants are commonly utilized during surgeries and to treat thrombotic diseases like stroke and deep vein thrombosis. However, conventional anticoagulants have serious side-effects, narrow therapeutic windows, and lack safe reversal agents (antidotes). Here, an alternative RNA origami displaying RNA aptamers as target-specific anticoagulant, is described. Improved design and construction techniques for self-folding, single-molecule RNA origami as a platform for displaying pre-selected RNA aptamers with precise orientational and spatial control, are reported. Nuclease resistance is added using 2’-fluoro-modified pyrimidines during *in vitro* transcription. When four aptamers are displayed on the RNA origami platform, the measured thrombin inhibition and anticoagulation activity is higher than observed for free aptamers, ssRNA-linked RNA aptamers, and RNA origami displaying fewer aptamers. Importantly, thrombin inhibition is immediately switched off by addition of specific reversal agents. Results for ssDNA and ssPNA (peptide nucleic acid) antidotes show restoration of 75% and 95% coagulation activity, respectively. To demonstrate potential for practical, long-term storage for clinical use, RNA origami was freeze-dried, and stored at room temperature. Freshly produced and freeze-dried RNA show identical levels of activity in coagulation assays. Compared to current commercial intravenous anticoagulants, RNA origami-based molecules show promise as safer alternatives with rapid activity switching for future therapeutic applications.

## 1. Introduction

The blood coagulation cascade is a complex enzymatic reaction network with many molecular components and intricate feedback control mechanisms. The ability to control and especially to down-regulate the coagulation cascade is necessary to human health in many situations. Anticoagulants have been used extensively in medical therapies to prevent and treat thromboembolism-related disorders including stroke, deep vein thrombosis, atrial fibrillation, and during cardiac bypass and percutaneous coronary intervention.^[1]^ Conventional drugs (e.g., warfarin and heparin) and novel anticoagulant drugs offer great benefits for medical uses; however, they are associated with serious side-effects, narrow therapeutic windows, and typically lack specific reversal agents.^[1d, 2]^ Therefore, innovative development of effective, new, target-specific anticoagulants with no severe side-effects and with specific antidotes is crucial to the health care enterprise.

Nucleic acid (NA) aptamers have been selected from libraries of randomized sequences to bind specifically with target molecules such as proteins, small molecules, inorganic materials, and whole cells.^[3]^ Due to their tunable molecular recognition capabilities, and their capacity for modular design, aptamers have been extensively developed in a wide range of applications including as biosensors, diagnostics, and therapeutics.^[3h, 4]^ Target-specific DNA and RNA aptamers that bind to enzymes of the coagulation cascade such as Factor IXa, Factor Xa, Factor XII/XIIa, and exosites-1 and −2 of thrombin, have been developed as anticoagulants and shown activity both *in vitro* and *in vivo*.^[5]^. Importantly, specific antidotes (i.e., reversal agents) for nucleic acid anticoagulants are automatically provided since nucleic acid strands with Watson-Crick complementary sequences will bind, unfold, and deactivate the aptamers.^[5b, 5f, 6]^ A short, Factor IXa-binding RNA aptamer anticoagulant was previously developed and tested in clinical trials.^[7]^ Because of its low molecular weight, the RNA was bulked-up by conjugation with a polyethylene glycol (PEG) group to avoid its being quickly removed from the blood via renal clearance. Unfortunately, some patients showed severe immune response to the coupled PEG moiety, therefore the Phase 3 clinical trial on the PEG-modified RNA anticoagulant was halted.^[8]^ The high molecular weight RNA origami platforms described here obviate the need for conjugation of any PEG-like bulking groups thus renewing interest in development of bare NA anticoagulants.

Nucleic acid nanotechnology provides a toolbox for designing programmable biomolecular platforms that can be decorated with functional NA motifs (ssDNA, ssRNA, aptamers, stem-loops, etc.), proteins (enzymes and antibodies), and other nanomaterials with nanoscale spatial control of placement and patterning.^[9]^ DNA and RNA nanostructures with arbitrary shapes and sizes have been constructed and developed.^[10]^ Due to their intrinsic properties, nucleic acid-based nanostructures have been used in both fundamental research and technological advances. NA nanostructures decorated with aptamers have been developed and utilized in a wide range of applications such as biosensing and therapeutics. ^[4, 9h, 11]^ For example, DNA weave tile nanostructures displaying multiple thrombin-binding DNA aptamers were shown to exhibit enhanced anticoagulation activity compared with free DNA aptamers.^[6b, 12]^ Reversal of anticoagulation activity was demonstrated by the addition of single-stranded DNA (ssDNA) antidotes. However, the DNA weave tiles were constructed from two strands of DNA thereby adding problems to large-scale production due to the need to balance stoichiometry and to purification issues related to removal of unwanted by-products. However, self-folding, single-molecule nucleic acid structures such as those described here can potentially overcome these problems.

RNA origami is a self-assembling nanostructure made of a single, self-folding molecule of RNA. Geary *et al.* reported co-transcriptional folding of RNA origami displaying kissing loops at the ends of RNA helices.^[10f]^ In the same paper, they demonstrated formation of hexagonal RNA lattices by interconnection of origami based on these kissing loop interactions. Furthermore, the self-assembly of single-stranded RNA origami of various shapes has been performed both *in vitro* and *in vivo*.^[10i, 13]^ RNA origami serves as a biomolecular platform with modularity, flexibility of design, and spatial control for displaying functional motifs, in particular RNA aptamers, with nanoscale precision. RNA origami has also been decorated with functional motifs such as fluorogenic RNA aptamers and demonstrated to perform as a novel biosensor inside cells using Förster resonance energy transfer (FRET).^[4]^ Additionally, protein-binding RNA aptamers were displayed on RNA origami constructs.^[9i]^

Recently, we reported unprecedented, single-strand RNA origami anticoagulants with significant *in vitro* activity.^[9h]^ The 2-helix RNA origami displaying two, thrombin-targeted RNA aptamers provided improved anticoagulation activity compared to free RNA aptamer and compared to DNA nanostructures decorated with thrombin-binding DNA aptamers. To enhance anticoagulation activity of RNA origami, we hypothesized that the number of RNA aptamers on each origami and the distances between exosite-1 and −2 RNA aptamers play important roles. Exosite-1 and 2 are anionic-binding domains located near the active sites of thrombin **(Figure 1A)**. In this work, we, therefore, develop new designs of single-stranded, 2-helix RNA origami (2HO-RNA) bearing four RNA aptamers. Additionally, we further developed and designed 3- and 4-helix RNA origami (3HO- and 4HO-RNA) that provide various distances between the aptamers. We kept the two aptamers on the same side of the origami and moved them apart by indexing their position on different helices as shown in Figure 2A-B. We chose well-studied exosite 1-binding aptamer called “RNA_R9D-14T_” and exosite 2-binding RNA aptamer called “Toggle-25t” for decoration on RNA origami (Figure 1).^[5c, 5f]^ To stabilize the RNA constructs under nuclease-containing conditions such as human plasma, 2’-fluoro-modified pyrimidine NTPs (2’F-dCTP and 2’F-dUTP) were incorporated in the nanostructures during *in vitro* transcription. We tested the specificity of binding of RNA origami against thrombin and other coagulation cascade proteins. The anticoagulation activity of the constructs was tested in activated partial thromboplastin time (aPTT) assays and with a confocal microscopy clot morphology assay. We also developed a practical, room-temperature storage method that may prove useful during future clinical use of these agents.

**Figure 1.**
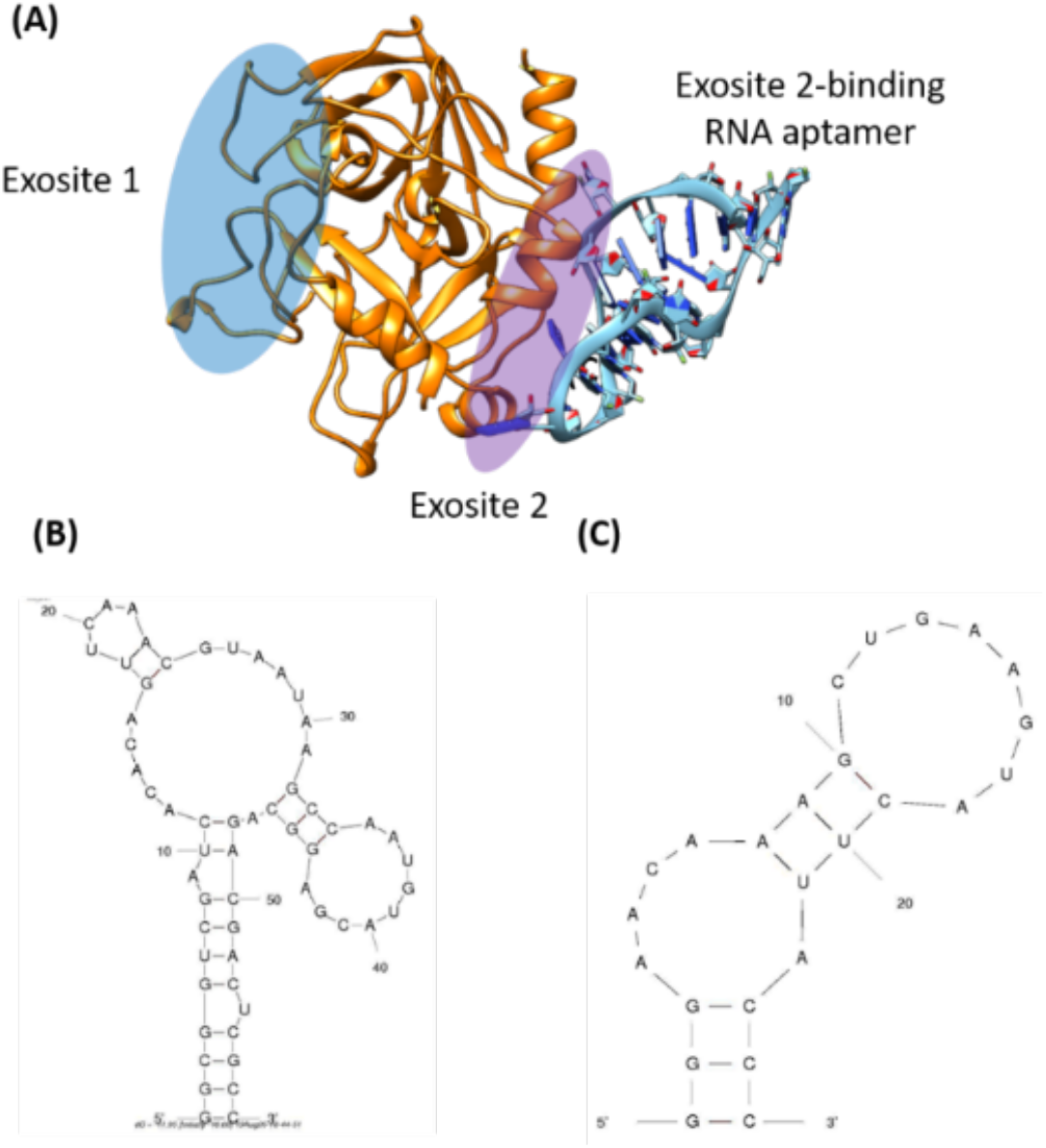
(A) Schematic drawing from the crystal structure of an exosite 2-binding RNA aptamer bound to thrombin (PDB: 3DD2).^[5c]^ Exosite 1 and 2 are highlighted in blue and purple, respectively. (B) Nucleotide sequence and secondary structures of exosite 1-, RNA_R9D-14T_, and (C) exosite 2-binding RNA aptamer, Toggle-25t. ^[5c, 5f]^ The secondary structures were computationally predicted by Mfold.^[16]^

**Figure 2.**
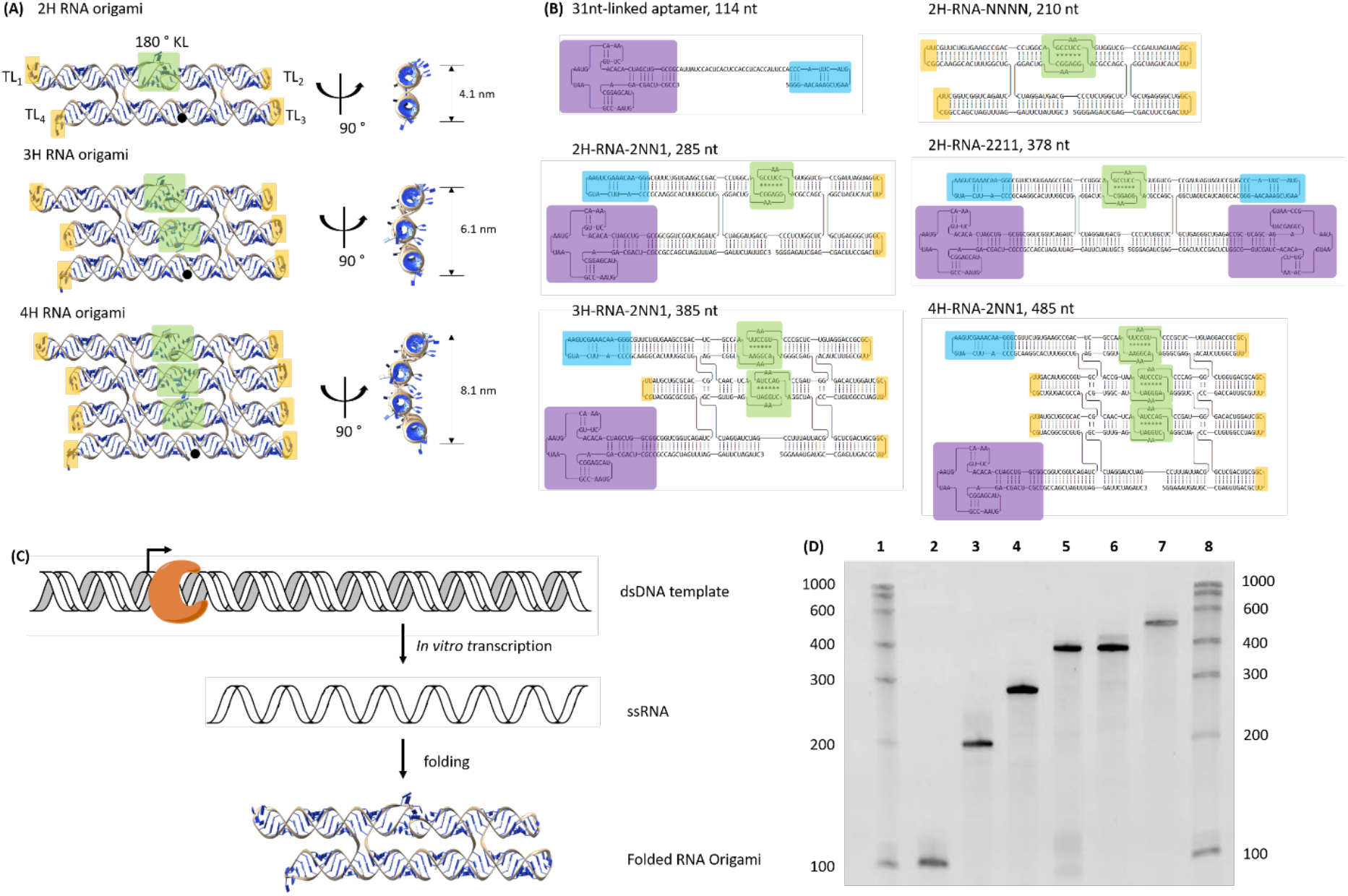
(A) 3D model of 2-, 3, and 4-helix RNA origami without thrombin aptamers (Front and side views). Black dots mark the 5’ ends of the origami molecules. (B) 2D schematics of various structures including ssRNA-linked aptamers, and RNA origami variants bearing zero, two, or four thrombin-binding aptamers at different positions. Exosite-1 and Exosite-2 binding aptamers are indicated in purple and blue rectangles, respectively. Tetra loops (TL) and 180° kissing loops (KL) are highlighted in yellow and green rectangles, respectively. Note that the variant naming conventions are described in the main text. (C) Schematic representation of *in vitro* transcription of RNA origami and heat-annealing (folding) process to form RNA origami. The orange crescent-like object is T7 RNA polymerase. (D) Characterization of RNA origami by denaturing polyacrylamide gel electrophoresis. Lane 1 and 8: ssRNA size markers, Lane 2: 31nt-linked aptamers (ss12), Lane 3: 2HF-NNNN, Lane 4: 2HF-2NN1, Lane 5: 2HF-2211, Lane 6: 3HF-2NN1, Lane 7: 4HF-2NN1.

## 2. Results and Discussion

### 2.1 RNA origami anticoagulant design

We adapted and modified 2H-,3H- and 4H-AE core RNA origami designs from previously described studies. ^[9h, 9i, 10f]^ Briefly, RNA origami structures comprise of double-helical domains, 180° kissing loops at the center of the structure, tetra-loops at the ends of helices and RNA crossover (strand exchange points) to link neighboring helices. 3D illustrations of RNA origami variants are shown in Figure 2A. To construct RNA origami anticoagulant, we fused thrombin-binding RNA aptamers in place of some of the tetraloops. The 2-helix RNA origami (2HO-RNA) contains four tetra loops at its four helical ends; these positions provide the four possible positions available for appending RNA aptamers. For naming RNA origami anticoagulants, we use a four-digit numbering system such as: 2HO-RNA-XXXX (Figure 2A). Each digit refers to the anchor position on RNA origami.

The exosite 1-(RNA_R9D-14T_) and exosite 2-binding RNA aptamer (Toggle-25t) are called “1” and “2”, respectively. Absence of RNA aptamer at some position is represented with “N”. For example, 2HO-RNA-2211 refers to 2-helix RNA origami displaying four RNA aptamers that contain two exosite 2-binding aptamers at positions 1 and 2 and two exosite 1-binding aptamers at positions 3 and 4. 2D illustrations of various RNA origami anticoagulants are depicted in Figure 2B.

### 2.2 Production of RNA origami anticoagulants

The production of long poly-ribonucleotides using chemical synthesis is limited by the efficiency of individual reactions. We produced long strands of RNA origami anticoagulants up to 485 nucleotides (nt) using *in vitro* transcription as shown in Figure 2C and detailed in the methods section. Synthesizing nuclease-resistant RNA anticoagulant is crucial since it will be used in RNase-containing environments such as human plasma and whole blood. The incorporation of 2’-fluoro-modified NTP into RNA aptamers and RNA origami provided stability against RNase.^[9h, 17]^ We, therefore, produced modified RNA origami by substitution of native CTP and UTP with 2’-fluoro-modified dCTP and dUTP as precursors for transcribed RNA origami. Transcripts containing 2’-F modifications were denoted with an ‘F’ in their name, while unmodified versions contain an ‘O’ in their name to indicate normal ribose sugars bearing 2’-OH moieties. To incorporate 2’F-dNTPs into RNA origami, mutant T7 RNA polymerase (Y639F) was used instead of native T7 RNA polymerase.^[18]^ Transcribed RNA origami anticoagulants were characterized using denaturing polyacrylamide gel electrophoresis. The modified RNA origami showed stability in RNase A (Figure S14). In this study, we tested six different RNA anticoagulant designs (Figure 2B and D).

### 2.3 Specific binding of RNA origami anticoagulants

We tested the binding of thrombin and other proteins to RNA origami bearing thrombin-binding RNA aptamers using a gel mobility shift assay. The self-assembled, 2’-fluoro-modified RNA origami constructs were incubated with thrombin, and the resulting complexes were characterized by native polyacrylamide gel electrophoresis (PAGE). As shown in Figure 3A, RNA nanostructures containing RNA aptamers (Fss12, 2HF-2NN1, 2HF-2211, 3HF-2NN1 and 4HF-2NN1) bind to thrombin in contrast to RNA origami without RNA aptamers (2HF-NNNN) which shows no band shift, indicating no interaction. We further examined the binding specificity of RNA origami by incubating with thrombin and three other proteins (Factor IXa and X from the coagulation cascade, and bovine serum albumin, BSA). The specific binding of RNA origami was characterized by native PAGE (Figure 3B, S8-13). The gel results showed that RNA origami bands migrate slower in the presence of thrombin (association is demonstrated). We further confirmed by performing both nucleic acid-staining and protein-staining on these gels. Our results indicate that RNA origami containing RNA aptamers specifically bound to thrombin and not to the other proteins tested.

**Figure 3.**
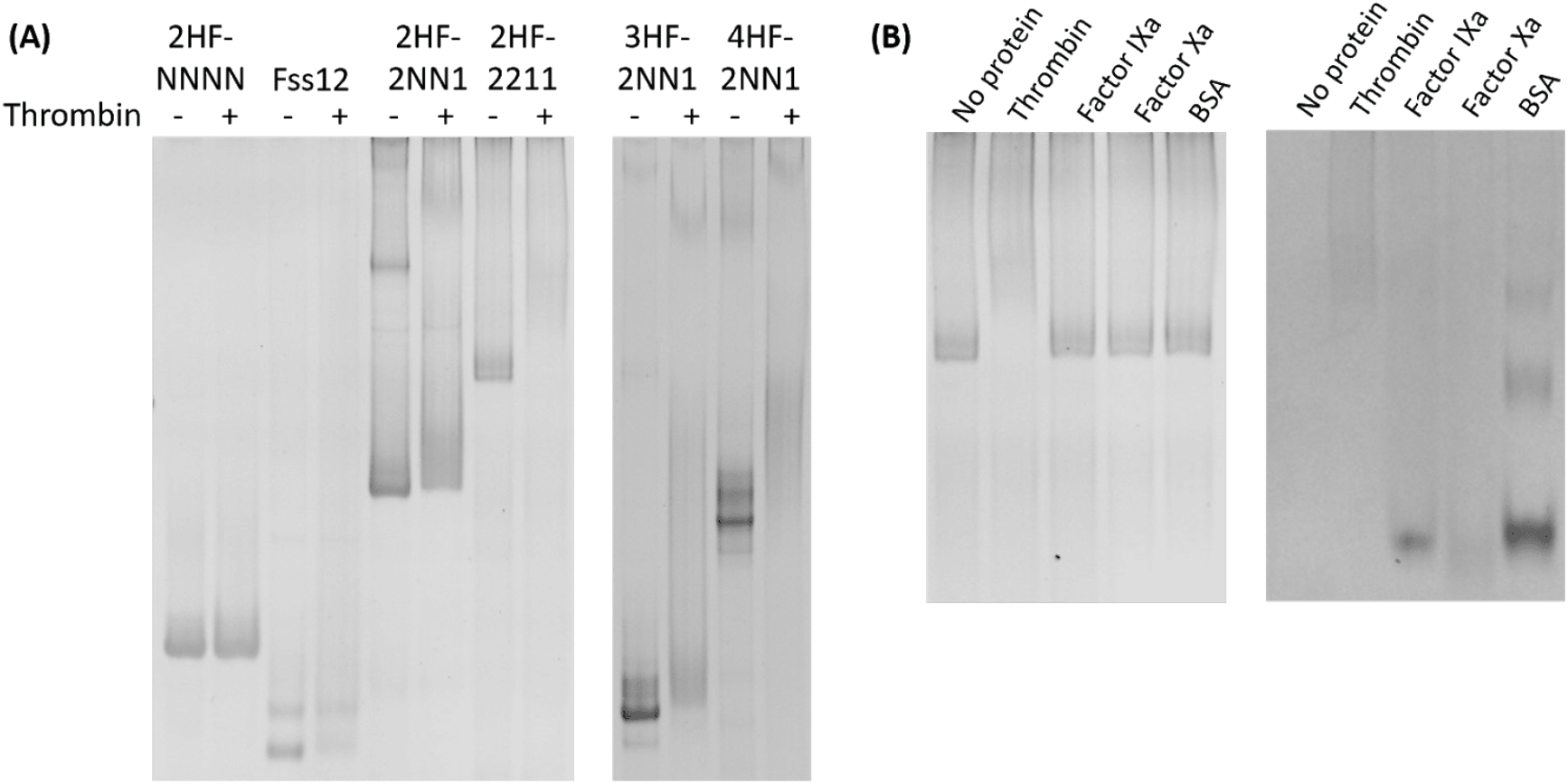
(A) Characterization of RNA origami bearing thrombin aptamers for binding with thrombin using 6% native polyacrylamide gel electrophoresis run at 150 V for 3 hr (left) and 6 hr (right). Gels were stained with ethidium bromide for nucleic acid staining. Lanes marked with − and + are the absence and presence of thrombin, respectively. (B) Specificity of binding of 2HF-2211 incubated with various proteins, including thrombin, Factor IXa, IXa and BSA. Left and right images are the same gel. Left: ethidium bromide-stained for nucleic acid. Right: Coomassie-stained for protein. As expected, RNA band shift is only observed with thrombin incubation.

### 2.4 Measured anticoagulation activity

The increasing local concentration of ligands due to designed clustering of aptamers on origami results in enhanced strength of binding interactions. To test a simpler model of local concentration effects, we designed a single-stranded RNA molecule with two thrombin-binding aptamers linked by a ssRNA tether (Fss12). We tested its anticoagulation activity using aPTT assay as described above. The clotting times of a mixed solution of two free aptamers and Fss12 are 47 and 74 seconds, respectively (Figure 4A). The anticoagulation activity of the ssRNA-linked RNA aptamers is around 1.5 times higher than the free aptamers. We further compared RNA origami-based anticoagulants (2HF-RNA-2NN1) with the Fss12 design where both molecules contain two RNA aptamers. Interestingly, the anticoagulation activity of 2HF-RNA-2NN1 is approx. 3 times higher than the Fss12. This suggests that the nanostructure of the RNA origami not only enhances binding by increasing local concentration, but also arranges the aptamers in specific positions that optimize thrombin inhibition. As expected, RNA origami without any aptamers (2HF-RNA-NNNN) showed no anticoagulant activity.

**Figure 4.**
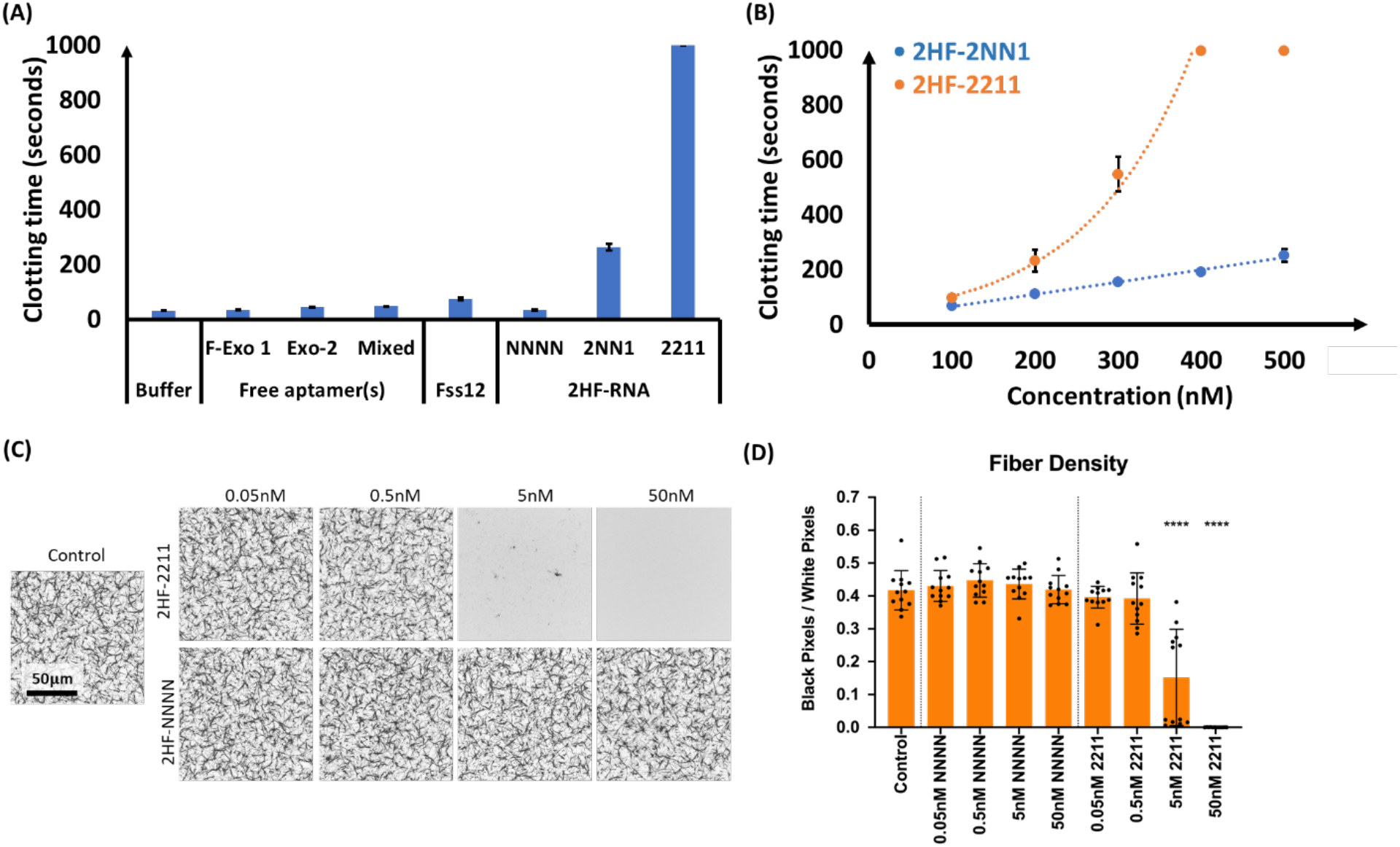
(A) Anticoagulation activity of RNA-based inhibitors using aPTT assay at 500 nM and (B) Concentration-dependent anticoagulant activity of the denoted RNA origami. (C) Representative concentration-dependent microscopic clot structure examination with RNA-based anticoagulants, 2HF-RNA-2211 (top) and 2HF-RNA-NNNN (negative control) (bottom), and positive control clot (left) using confocal microscopy. (D) Quantification of fiber density (black pixels representing fiber and white pixels representing background) from confocal microscopy images. Mean +/− standard deviations are shown. ****p<0.0001.

The 2-helix RNA origami design offers 4 positions for aptamer placement. To enhance the anticoagulant effect of the RNA origami, we created a 2-helix origami displaying four RNA aptamers, two for exosite 1 and two for exosite 2, named 2HF-RNA-2211 (Figure 2B). Activity of the RNA origami was tested using aPTT coagulation assays. As shown in Figure 4A, the average clotting times of 2HF-RNA-2NN1 at 500 nM were approximately 260 seconds, while the clot time for 2HF-RNA-2211 at 400 and 500 nM reached the maximum limit of the coagulometer at 999 seconds. Surprisingly, the clotting time of 2HF-RNA-2211 (500 nM) was more than double the clotting time of 500 nM 2HF-RNA-2NN1. This suggests a non-linear relationship between appended aptamer number and clot time. To better understand the relationships between the 2-aptamer and 4-aptamer designs, the coagulation activity of 2HF-RNA-2NN1 and 2HF-RNA-2211 was compared at concentrations ranging from 100 nM to 500 nM in pooled human plasma. As seen in Figure 4B, the 2-aptamer design displayed a linear relationship between concentration and clotting time. Surprisingly, the 4-aptamer design showed a super-linear relationship between clotting time and concentration. The mechanism for this behavior is unknown and will be investigated further.

Anticoagulant activity of 2HF-RNA-2211 was also observed in confocal microscopy assays examining clot structure. Control clots, made with purified fibrinogen and thrombin, formed robust clot networks as seen in Figure 4C, with fibrin fiber densities of 0.42 +/− 0.06 (mean +/− standard deviation). Clots formed with 2HF-RNA-NNNN did not significantly alter clot structure or fiber density at any concentration tested compared to control clots. However, upon the addition of 2HF-RNA-2211, clot structure was significantly impaired at 5 nM and 50 nM concentrations with significant decreases in fiber density, as shown in Figure 4D. At 5 nM 2HF-RNA-2211, fiber density is significantly lower than control clots or clots formed with the same molecule at lower concentrations (fiber density = 0.15 +/− 0.1, p<0.0001). At 50 nM 2HF-RNA-2211, no fibrin fibers were observed. The complete lack of clot microstructure indicates significant anticoagulant activity and complete inhibition of fibrin polymerization.

To test the effect of spatial positioning of the aptamers, we designed three RNA origami bearing two RNA aptamers and 2-, 3-, and 4- helix origami structures (2HF-2NN1, 3HF-2NN1, 4HF-2NN1, respectively) as shown in Figure 2B. The estimated width of 2HF−, 3HF−, and 4HF-RNA origami measured from 3D models are approximately 4, 6, and 8 nm, respectively. However, the actual distance between two RNA aptamers on an origami may be somewhat different due to the inherent flexibility of these structures. Based on X-ray crystallography and electron microscopic analysis, it is known that thrombin is a globular protein with diameter of approximately 5 nm.^[19]^ The X-ray crystallographic data shows that the structure is 4.5 × 4.5 × 5.0 nm^3^ and determine the distance between exosite 1 and 2 of 4.5 – 5.0 nm.^[19b]^ We tested the anticoagulant activity of three designs of RNA origami bearing two RNA aptamers using aPTT assay at 1 uM. The activity of 3HF-2NN1 and 4HF-2NN1 were slightly higher than 2HF-2NN1 but not significantly so (Figure 5). Since it is more difficult and expensive to produce 3HF- and 4HF-2NN1 than the smaller two-helix variant, and since their activity was not significantly higher, we continued further experiments with 2HF-RNA origami.

**Figure 5.**
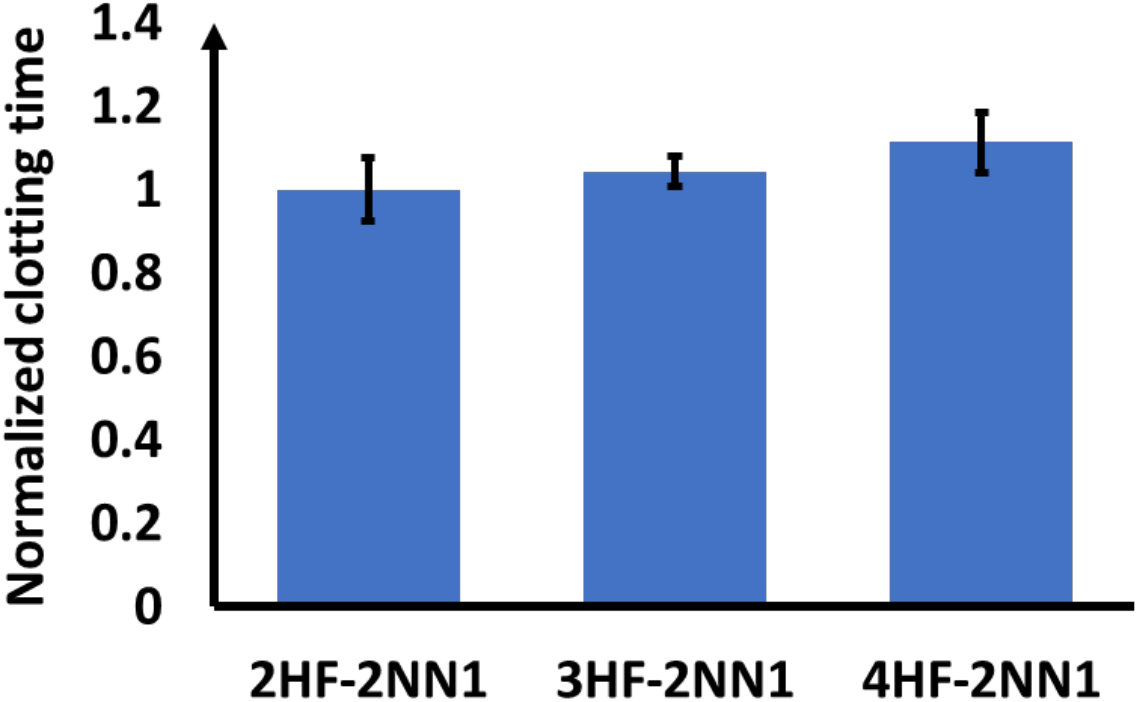
Anticoagulation activities of 2-helix, 3-helix, and 4-helix RNA origami bearing two thrombin aptamers. Clotting times were measured using aPTT assay. Mean of normalized clotting times +/− standard deviations are shown.

### 2.5 Practical usage of RNA origami and long-term storage

The storage of RNA can be difficult due to its chemical and enzymatic instability. Previously, our chemically-modified RNA origami structures have been shown to last for up to 3 months in solution in a folding buffer at 4 °C.^[9h]^ Although this is potentially promising for clinical use, room temperature storage would be more advantageous and reduce storage costs. Storing RNA in a dry state, rather than in buffer, increases its resistance to degradation at room temperature. To test this concept, we compared the anticoagulation activity of freshly produced 2HF-RNA-2NN1 with freeze-dried 2HF-RNA-2NN1 using aPTT coagulation assays. The RNA samples were folded prior to lyophilization and stored at room temperature overnight. The origami was resuspended in water and immediately tested at a concentration of 1 μM. As shown in Figure 6, it was found that the anticoagulation activity remained consistent following the freeze-drying process, as clotting time was approximately 70 seconds both prior to and after lyophilization. This suggests that our RNA origami is not negatively affected by the freeze-drying process, and can be reliably stored at room temperature, providing benefits for future clinical use.

**Figure 6.**
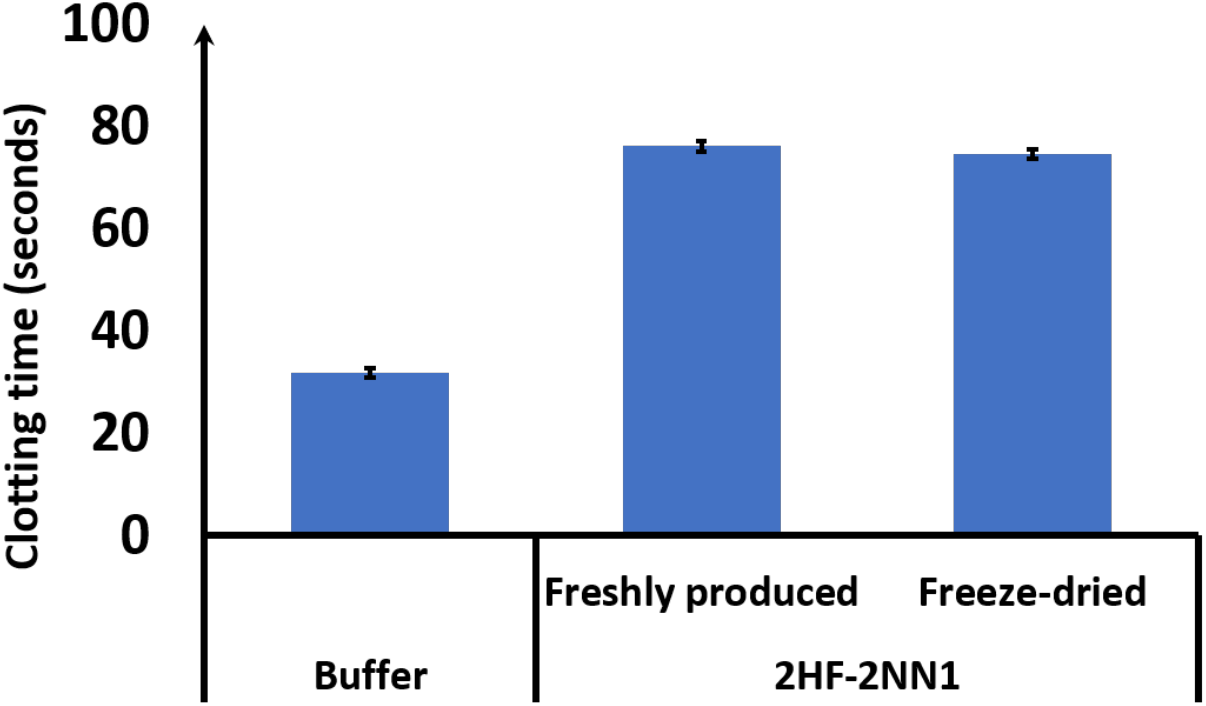
Stability of RNA origami after freeze-drying. The anticoagulation activity of freshly produced 2HF-2NN1 (1 μM) and resuspended 2HF-2NN1 after the freeze-drying process. Mean +/− standard deviation.

### 2.6 Reversible thrombin inhibition: specific ssDNA and ssPNA antidotes

The ability to reverse the anticoagulation activity of active anticoagulant with a specific antidote would be very helpful in a range of medical procedures. Nucleic acid-based anticoagulants offer excellent benefits over chemical-based anticoagulants by providing rapid, specific reversal agent in the form of strands with sequence complementary to that of the aptamer. Specific antidotes allow us to reverse thrombin inhibition with potential benefits to medical and therapeutic uses. We designed two nucleic acid antidotes, complementary to exosite-1- and exosite-2-binding RNA aptamers (Figure 7A). The reversal agent is a complementary sequence of RNA aptamer that hybridizes and unfolds RNA aptamer. Antidotes for exosite-1 and −2 aptamers are 20 and 19 nucleotides long, respectively, and are complementary to the RNA aptamer sequences. Reversal of the anticoagulant activity was tested by aPTT assay. We found that recovery of coagulation activity was 75% by the addition of a specific DNA antidote (Aptamers to antidote ratio is 1:9) (Figure. 7B). For therapeutic purposes, highly effective and specific antidote allowing physicians to switch on and off anticoagulation activity would be highly beneficial. Therefore, we designed and developed higher performance reversal agents using PNA antidotes. To improve the efficiency of antidote, elimination of electrostatic repulsion between phosphate backbone of RNA aptamer and its antidote plays a crucial role. Peptide nucleic acid is a non-natural nucleic acid that has no charge of its backbone and results in strong interaction between PNA/DNA and PNA/RNA complexes. Due to the high affinity of PNA binding with DNA and RNA, PNA has been employed for blocking of transcription by binding with genomic DNA and inhibiting translation by strongly binding mRNA.^[20]^ Therefore, we further developed and tested high efficiency, specific PNA antidotes. We observed a recovery of 95% of thrombin activity thereby outperforming the DNA antidotes. The combination of RNA origami anticoagulant and its specific PNA antidote is highly effective, offering many benefits to medical therapeutic uses over conventional anticoagulants that lack specific antidotes.

**Figure 7.**
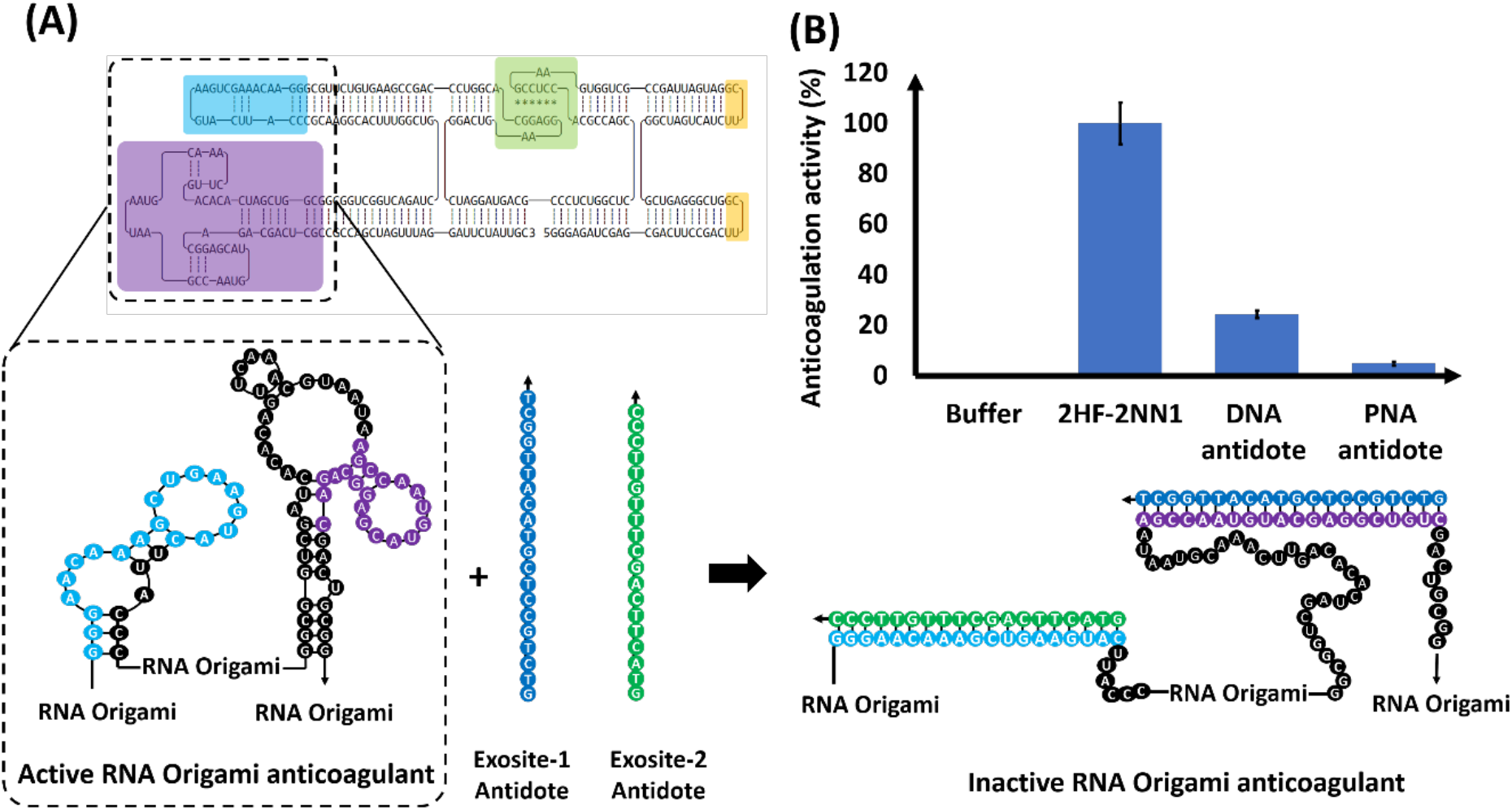
Reversibility of RNA origami anticoagulation via ssDNA and ssPNA antidotes. (A) 2D illustration of active and inactive 2HF-2NN1 anticoagulant. Exosite-1 and 2 aptamers are highlighted in purple and blue, respectively. (B) Recovery of thrombin activity by addition of specific DNA and PNA antidotes. Percent activity remaining +/− standard deviation.

## 3. Conclusion

We designed and constructed nuclease-resistant, self-folding, single-molecule RNA origami platforms decorated with multiple thrombin-binding RNA aptamers and have shown them to be highly effective anticoagulants for direct inhibition of thrombin. We also developed specific antidotes or reversal agents for these anticoagulants. The 2-, 3-, and 4- helix RNA origami constructs were synthesized by *in vitro* transcription. The 4-helix design offers up to eight possible helix ends as positions to display RNA aptamers with precise spatial control. These aptamer-on-origami molecules bind specifically to thrombin and inhibit coagulation more effectively than free aptamer or ssRNA-linked aptamers. The variant with four aptamers on 2-helix RNA origami offers a significant enhancement of anticoagulation activity compared with versions bearing two aptamers. Thrombin inhibition can be reversed and clotting activity recovered by addition of specific antidotes. A PNA antidote, with no net charge, offers a highly effective antidote with a recovery efficiency of 95 %. Importantly, the RNA origami anticoagulant is stable and retains its activity after undergoing freeze-drying and re-dissolution - a great benefit to long-term storage and clinical use. These RNA origami-based anticoagulants have high molecular weight, so no further chemical modification will be necessary to increase their size for avoiding kidney clearance in *in vivo* studies. This RNA origami technology provides a novel biomolecular toolbox that could potentially be utilized in a variety of fields, including surgical and therapeutic uses.

## 4. Experimental section

### Materials

Phusion High-Fidelity DNA polymerase, deoxynucleotide triphosphates (dNTPs), nucleotide triphosphates (NTPs), DNase I, RNase A, and nuclease-free water were purchased from ThermoFisher Scientific. Bovine serum albumin (BSA) was purchased from Sigma-Aldrich. Thrombin, Factor IXa, and Factor Xa were purchased from Haematologic Technologies, Inc. The Monarch RNA clean-up kit and NEBbuilder HiFi DNA assembly master mix were purchased from New England Biolabs. Custom oligonucleotides and custom G-block DNA were purchased from Integrated DNA Technologies (IDT). DNA primers were ordered desalted from IDT and used without further purification. 2’-Fluoro-2’-deoxycytidine-5’-Triphosphate (2’F-dCTP) and 2’-Fluoro-2’-deoxyuridine-5’-Triphosphate (2’F-dUTP) were purchased from Trilink Biotechnologies, LLC. Mutant T7 polymerase Y639F was produced in our laboratory. Pooled human plasma was purchased from George King Bio-Medical, Inc. Single-stranded PNA antidotes were ordered with HPLC purification from PNA Bio. CaCl_2_ solution and aPTT reagent were purchased from Trinity Biotech Plc. GFX PCR purification kit was purchased from GE Healthcare.

### RNA origami design

Design of RNA origami constructs has previously been described by Sparvath *et al*.^[14]^ We adapted one of the core RNA origami structures from previous studies.^[9h, 9i, 10f]^ Briefly, we first obtained coordinates and models for 3D structures of assembled RNA origami from various resources such as double-stranded A-form RNA, 180° kissing loop, tetraloop and exosite-2 binding RNA aptamer. The 3D crystallographic structure of exosite-1 binding aptamers has not been solved. First, a double-stranded A-form RNA was constructed by make-rna server online website (http://structure.usc.edu/makena/server.html). The 180° kissing loop was obtained from the RNA junction website (https://rnajunc-tion.ncifcrf.gov). The tetraloop (PDB: 1F7Y) and exosite 2-binding RNA aptamer (PDB: 3DD2) were obtained from the RCSB PDB website (www.rcsb.org/). Prior to model assembly, the tetraloop and RNA aptamer were extracted from a larger structure using Swiss-PdbViewer (https://spdbv.vital-it.ch). Next, all components were aligned with UCSF Chimera software (https://www.cgl.ucsf.edu/chimera/) in order to create a 3D model of RNA origami. The aligned RNA origami structure was converted to a single-stranded RNA origami by using Perl script called “ligate.pl” that can be downloaded from www.andersen-lab.dk. The ligated single-stranded RNA structure was converted to a 2D model using Assemble2 software (https://bioinformatics.org/S2S/).

For RNA origami sequence design, we followed RNA origami design tutorial published by Sparvath et al.^[14]^ The 2D ribbon models of RNA origami were converted to NUPACK code by using a Perl script (download at www.andersen-lab.dk). The output code was pasted into NUPACK, online software (www.nupack.org) to generate RNA sequences.^[15]^ The sequence resulting from this process was manually edited. To predict RNA folding, the RNA sequences were tested with two computational RNA folding software packages, NUPACK and Mfold (http://unafold.rna.albany.edu/?q=mfold).^[15–16]^ The 2D ribbon models and computational analysis of RNA origami folding are shown in supporting information (Figure S1-6). Kissing loop interactions are not shown in folding predictions from online software. The results from computational analysis agreed with 2D ribbon designs as shown in Figure S1-6. The RNA sequence predicted to fold to the desired structure was converted to its DNA template form. Finally, the T7 RNA polymerase promoter sequence was added to the 5’end of DNA sequence and the template was ordered via custom oligonucleotide synthesis (IDT).

### Amplification of DNA template

DNA template was amplified from double stranded DNA G-block (IDT) using polymerase chain reaction (PCR). The PCR reaction consisted of 1x HF reaction buffer (ThermoFisher Scientific), 0.5 uM each for forward and reverse primers (IDT), 0.2 mM dNTP, 4 nM DNA template, and 0.02 unit/μL Phusion High-Fidelity DNA polymerase (ThermoFisher Scientific). Nuclease-free water was added to adjust to the desired volume, typically 50 or 100 mL. The PCR reaction was incubated as follows: (1) denaturing at 98 °C for 10 seconds, (2) annealing at 63 °C for 15 seconds, (3) extension at 72 °C for 20 seconds. These steps were repeated for 30 cycles. Following amplification, samples were characterized using agarose gel electrophoresis and purified using GFX DNA purification kit (GE Healthcare).

The 2H-2211 DNA template design could not be produced as a single G-block sequence due to repeated subsequences. Thus, the 2H-2211 DNA design was ordered as two separate G-block templates, referred to as fragment 1 and fragment 2. Prior to the amplification steps mentioned above, fragments 1 and 2 were joined using Gibson Assembly. Gibson Assembly was performed using NEBuilder HiFi DNA Assembly Kit. For assembly mixture, 0.1 pmol of each DNA fragment was added to 10 μL of NEBuilder HiFi DNA Master Mix. Nuclease-free water was added to adjust the volume to 20 μL. The mixture was incubated in a thermocycler at 50 °C for 15 minutes. For DNA amplification, 10 μL of Gibson Assembly reaction mixture was used in place of G-block template in the PCR reaction mixture. All other step details and component concentrations were according to the instructions above.

### Production and folding of RNA origami anticoagulants

RNA origami molecules were produced by *in vitro* transcription. The transcription reaction contained 5 ng/μL purified DNA template, 5 mM freshly prepared DTT, 2.5 mM each of NTPs (2’-fluoro-modified dCTP, - dUTP, ATP and GTP), 100 μg/mL mutant T7 polymerase Y639F, made-in-house. The reaction was transcribed at 37 °C for 16 hours. The transcribed RNA origami was purified using Monarch RNA clean-up kit. After purification, the concentration of purified RNA origami was measured by UV-spectrophotometry with absorbance at 260 nm (Nanodrop 3000C spectrophotometer, ThermoFisher Scientific). To fold RNA origami, the purified RNA product was diluted in 1x annealing buffer (20 mM HEPES pH 7.4, 150 mM NaCl, and 2 mM CaCl_2_) to desired concentration. The RNA product was heated at 95 °C for 5 minutes and let set on a benchtop to anneal/cool for 30 minutes.

### Specific binding of RNA origami with thrombin

The folded RNA origami (5 pmol) diluted in 1x annealing buffer was incubated with various proteins at 37 °C for 1 hr. The RNA origami-protein samples were characterized by 6% native polyacrylamide gel electrophoresis. After running, the gel was stained with ethidium bromide for nucleic acid visualized by UV imager (ProteinSimple Imager). Subsequently, the same gel was further stained with Coomassie blue for protein visualization and de-stained by soaking in Milli-Q water. The gel was imaged with white light in a ProteinSimple Imager.

### Anticoagulation activity of RNA origami by aPTT assay and confocal miscroscopy clot morphology assay

To test the anticoagulation activity of the RNA origami, an ST4 Coagulometer (Diagnostica Stago) was used to perform aPTT coagulation assays. Prior to testing, aPTT reagent was dissolved using 5 mL of ultrapure water and set at room temperature for 30 minutes. The coagulometer was set to aPTT and a temperature of 37 °C. To begin the test, a plastic cuvette (Diagnostica Stago) was placed into position and a magnetic bead (Diagnostica Stago) was placed into each cuvette well. 50 μL of pooled human blood plasma (George King Bio-Medical, Inc.) and 50 μL of aPTT reagent were pipetted into each cuvette well and allowed to incubate for 300 seconds. RNA origami (16.67 μL) was pipetted into each cuvette well and allowed to further incubate for 300 seconds. The cuvette was placed into the rightmost slot and the pipette key was pressed to initiate magnetic bead movement. Fifty microliters of CaCl_2_ was pipetted into the first well and the clot timer was immediately started using the pipette trigger. This process was repeated for each well and the clot times were recorded.

To further test the anticoagulation activity, confocal microscopy was utilized to examine fibrin clot structure. Fifty microliter clots were formed from purified fibrinogen using 0.25 U/mL human alpha-thrombin (Fisher Scientific), 2 mg/mL fibrinogen (Enzyme Research Laboratories), 0.1 mg/mL Alexa-Fluor 488 labeled fibrinogen for visualization (Thermo Fisher Scientific), all in HEPES buffer (25 mM HEPES, 150 mM NaCl, 5 mM CaCl_2_, pH 7.4). RNA (2HF-2211 or 2HF-NNNN) was incorporated at various concentrations (0.05, 0.5, 5, 50 nM), diluted in HEPES buffer (20 mM HEPES, 150 mM NaCl, 2 mM CaCl_2_, pH 7.4). For each condition, two duplicate clots were analyzed. Clots were formed between a glass microscope slide and a coverslip and allowed to polymerize for a minimum of three hours prior to imaging on a Zeiss Laser Scanning Microscope (LSM 710, Zeiss Inc., White Plains, NY, USA). A z-stack was imaged at a thickness of 1.89 μm at a 63x objective magnification. Three z-stack images were taken per duplicate. Image analysis was performed using ImageJ software to make 8-bit 3D projections of z-stack images. Fiber density was quantified by measuring the ratio of black (fibers) to white (background) pixels in each binary image.

### Room-temperature storage, freeze-dried RNA origami

The folded RNA origami in annealing buffer was frozen and then freeze-dried using a standard lyophilizer. The dried RNA origami was stored at room temperature until use. Prior to use, the dried RNA origami was re-dissolved in nuclease-free water. The anticoagulation activity of RNA origami was tested by aPTT assay.

### Reversal of anticoagulation activity with ssDNA and ssPNA antidotes

Anticoagulant activity of 2HF-2NN1 was reversed using ssDNA purchased from IDT and ssPNA (peptide-nucleic-acid) antidote strands purchased from PNA Bio. A solution containing two specific antidotes (500 μM, 1.5 μL for exosite-1 antidote and 500 μM, 1.5 μL of exosite-2 antidote with aptamer: antidote ratio approximately 1:9) were heated up to 90 °C for 10 min prior to use. To study the effect of antidote, aPTT coagulation assays were performed as described above. However, after the addition of RNA origami and the 300 seconds incubation, 3 μL of DNA or PNA antidote solution was added and incubated for an additional 300 seconds. For the control sample with no antidote, 3 μL of annealing buffer was added in place of the antidote solution. Clotting times were recorded for further analysis.

## Supporting Information

Supporting Information is available following this preprint text.

## Acknowledgements

This project was supported by NSF grants: IRES-1559077, BMAT-1709010, and BME-1603179. The authors also acknowledge support from American Heart Association Predoctoral Fellowship (18PRE33990338) to EM. Dr. Abhichart Krissanaprasit and Carson Key contributed equally to this work.

