## Supplemental Information for "Enhanced Anticoagulation Activity of Functional RNA Origami"

**RNA sequence of 2H-NNNN, 210 nt**

GGGAGAUCGAGCGACUUCCGACUUCGGUCGGGAGUCGGGCUAGUCAUCUUCGG

AUGAUUAGCCGCUGGUGAAGCCUCCACGCCAGCCUCGGUCUCCCGCAGUAGGA

UCGGACUGAAGGAGGCACGGUCCCAGCCGAAGUGUCUUGCUUCGGCAAGGCAC

UUUGGCUGCUAGACUGGCUGGCUUCGGCCAGCUAGUUUAGGAUUCUAUUGC

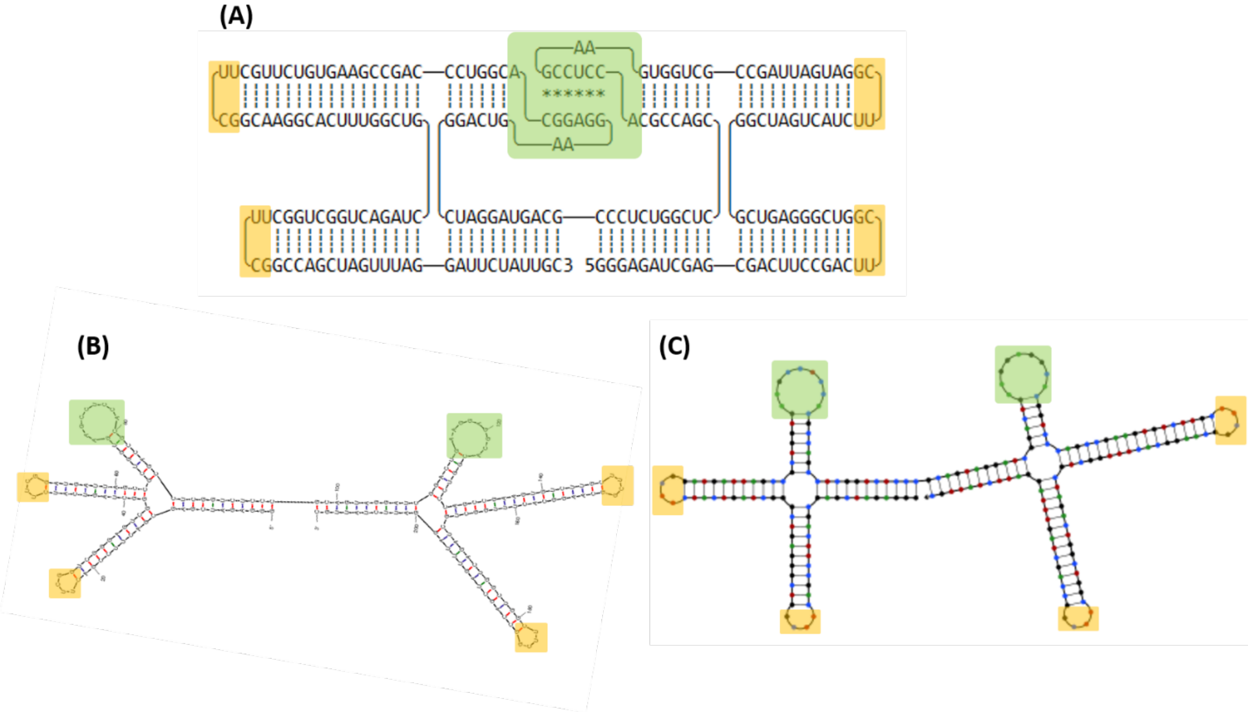


**Figure S1** (A) 2D ribbon model of 2H-NNNN RNA origami, without thrombin-binding aptamers that used for negative control. (B and C) Computational analysis of RNA origami folding analyzed by Mfold RNA and NUPACK, online software. Yellow and green rectangles indicate tetra loops and kissing loop motifs, respectively. The KL motifs were not shown in NUPACK and Mfold results.

**RNA sequence of 2H-2NN1, 285 nt**

GGGAGAUCGAGCGACUUCCGACUUCGGUCGGGAGUCGGGCUAGUCAUCUUCGG

AUGAUUAGCCGCUGGUGAAGCCUCCACGCCAGCCUCGGUCUCCCGCAGUAGGA

UCGGACUGAAGGAGGCACGGUCCCAGCCGAAGUGUCUUGCGGGAACAAAGCUG

AAGUACUUACCCGCAAGGCACUUUGGCUGCUAGACUGGCUGGCGGCGGUCGAU

CACACAGUUCAAACGUAAUAAGCCAAUGUACGAGGCAGACGACUCGCCGCCAG

CUAGUUUAGGAUUCUAUUGC

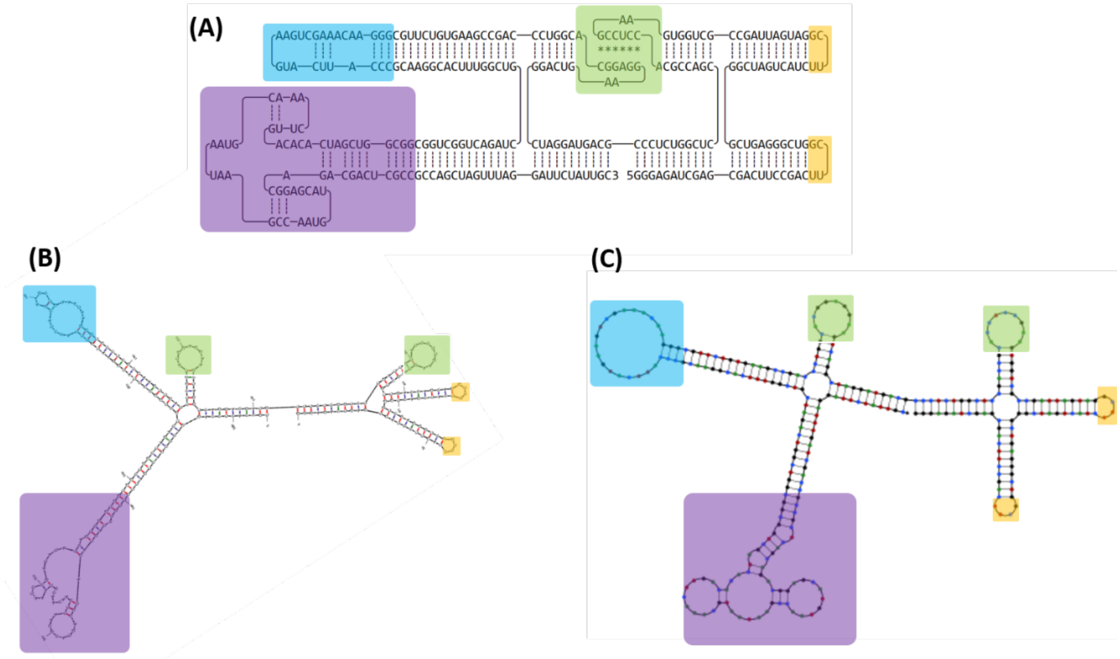


**Figure S2** (A) 2D ribbon model of 2H-2NN1 RNA origami anticoagulant. (B and C) Computational analysis of RNA origami folding analyzed by Mfold RNA and NUPACK, online software. Blue and purple represent exosite-I, and exosite-II-binding aptamers, respectively. Yellow and green rectangles indicate tetra loops and kissing loop motifs. The KL motifs were not shown in NUPACK and Mfold results.

**RNA sequence of 2H-2211, 378 nt**

GGGAGAUCGAGCGACUUCCGACUCUGGCGGUCGAUCACACAGUUCAAACGUAA

UAAGCCAAUGUACGAGGCAGACGACUCGCCAGAGUCGGGAGUCGGGCUAGUCA

UCAGGCACGGGAACAAAGCUGAAGUACUUACCCGUGCCUGAUGAUUAGCCGCU

GGUGAAGCCUCCACGCCAGCCUCGGUCUCCCGCAGUAGGAUCGGACUGAAGGA

GGCACGGUCCCAGCCGAAGUGUCUUGCGGGAACAAAGCUGAAGUACUUACCCG

CAAGGCACUUUGGCUGCUAGACUGGCUGGCGGCGGUCGAUCACACAGUUCAAA

CGUAAUAAGCCAAUGUACGAGGCAGACGACUCGCCGCCAGCUAGUUUAGGAUU

CUAUUGC

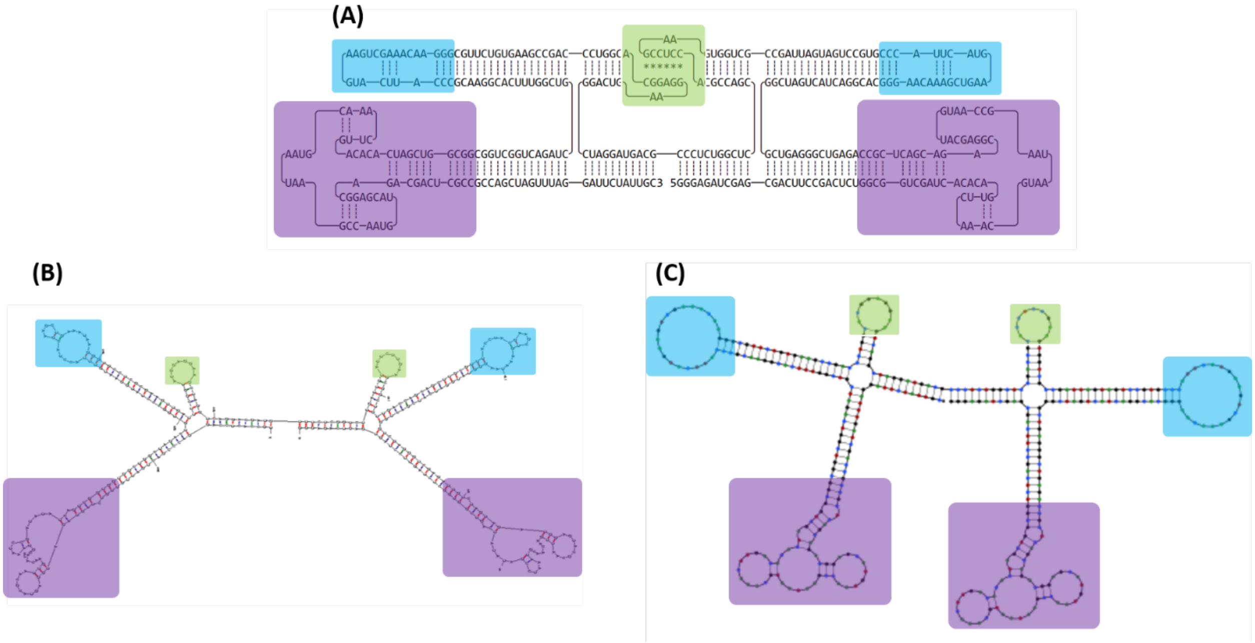


**Figure S3** (A) 2D illustration of 2H-2211 RNA origami anticoagulant. (B and C) Computational analysis of RNA origami folding analyzed by Mfold RNA and NUPACK, online software. Blue and purple rectangles represent exosite-I, and exosite-II-binding aptamers, respectively. Yellow and green rectangles indicate tetra loops and kissing loop motifs.

**RNA sequence of 31nt-linked aptamers, 114 nt**

GGGAACAAAGCUGAAGUACUUACCCACCUUACCACUCCACCUCACUCACCUAU

UACGGCGGUCGAUCACACAGUUCAAACGUAAUAAGCCAAUGUACGAGGCAGAC

GACUCGCC

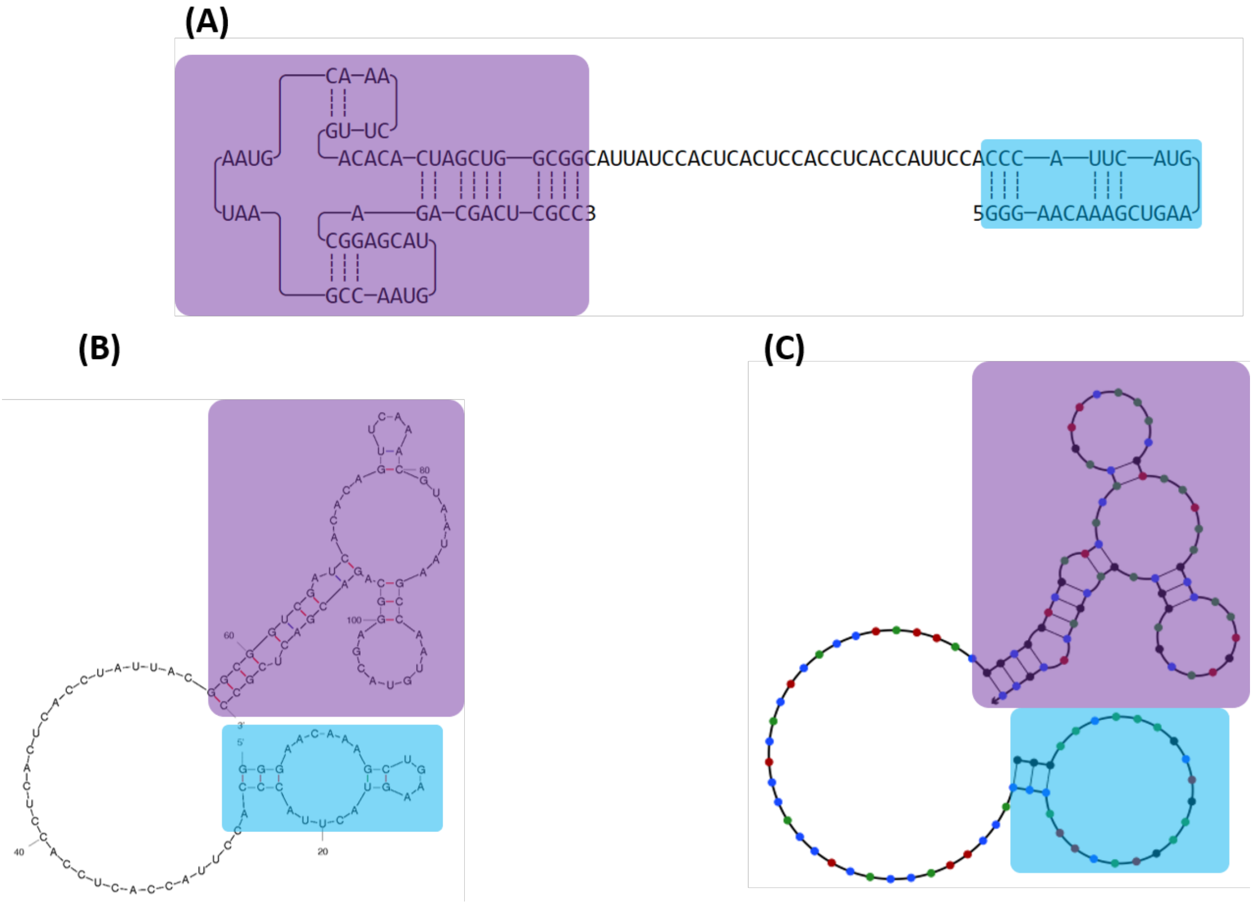


**Figure S4** (A) 2D illustration of 31nt-linked two thrombin aptamers. (B and C) Computational analysis of RNA origami folding analyzed by Mfold RNA and NUPACK, online software. Blue and purple rectangles represent exosite-I, and exosite-II-binding aptamers, respectively.

**RNA sequence of 3H-2NN1, 385 nt**

GGAAAUGAUGCCGAGUUGACGCUUCGGCGUCAGCUCGCCCUGUGGCCUAGUUC
GCUAGGUCACAGACAUCUUGGCGUUCGCGCCAGGAUGUCUCGCCCAAUUCCGU
AGGGCGAGGGUAGCCAAAUCCAGAGGCUAGCAUUAUUUCCGAUCUAGGAUCGC
GUUGAGAACUGGAUACUCAACAGCGGUAAACGGAAAACCGCUCAGCCGAAGUG
UCUUGCGGGAACAAAGCUGAAGUACUUACCCGCAAGGCACUUUGGCUGGCCAC
GCGUCGUAUUCGUACGGCGCGUGCUAGACUGGCUGGCGGCGGUCGAUCACACA
GUUCAAACGUAAUAAGCCAAUGUACGAGGCAGACGACUCGCCGCCAGCUAGUU
UAGGAUUCUAGAUC

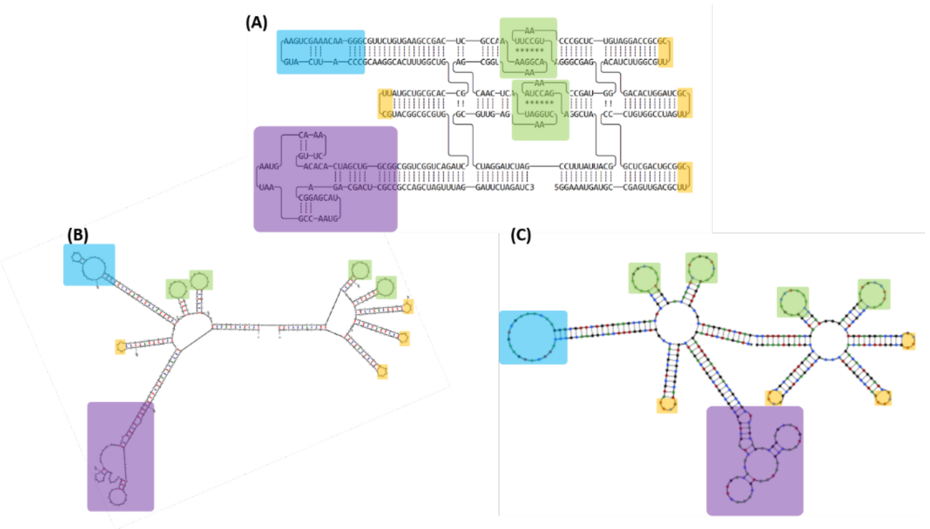


**Figure S5** (A) 2D illustration of 3H-2NN1 RNA origami anticoagulant. (B and C) Computational analysis of RNA origami folding analyzed by Mfold RNA and NUPACK, online software. Blue and purple rectangles represent exosite-I, and exosite-II-binding aptamers, respectively. Yellow and green rectangles indicate tetra loops and kissing loop motifs.

**RNA sequence of 4H-2NN1, 485 nt**

GGAAAUGAUGCCGAGUUGACGCUUCGGCGUCAGCUCGCCCUGUGGCCUAGUUC

GCUAGGUCACAGCCGACCAUUGCGUUUCGACGCAGUGGUCACAUCUUGGCGUU

CGCGCCAGGAUGUCUCGCCCAAUUCCGUAGGGCGAGGGGACCCAAAUCCCUAG

GGUCGGUAGCCAAAUCCAGAGGCUAGCAUUAUUUCCGAUCUAGGAUCGCGUUG

AGAACUGGAUACUCAACCGUGGCAUAAAGGGAUAAUGCCAAGCGGUAAACGGA

AAACCGCUCAGCCGAAGUGUCUUGCGGGAACAAAGCUGAAGUACUUACCCGCA

AGGCACUUUGGCUGCGUGGCGUUACAGUUCGCUGUGACGCCAGCCACGCGUCG

UAUUCGUACGGCGCGUGCUAGACUGGCUGGCGGCGGUCGAUCACACAGUUCAA

ACGUAAUAAGCCAAUGUACGAGGCAGACGACUCGCCGCCAGCUAGUUUAGGAU

UCUAGAUC

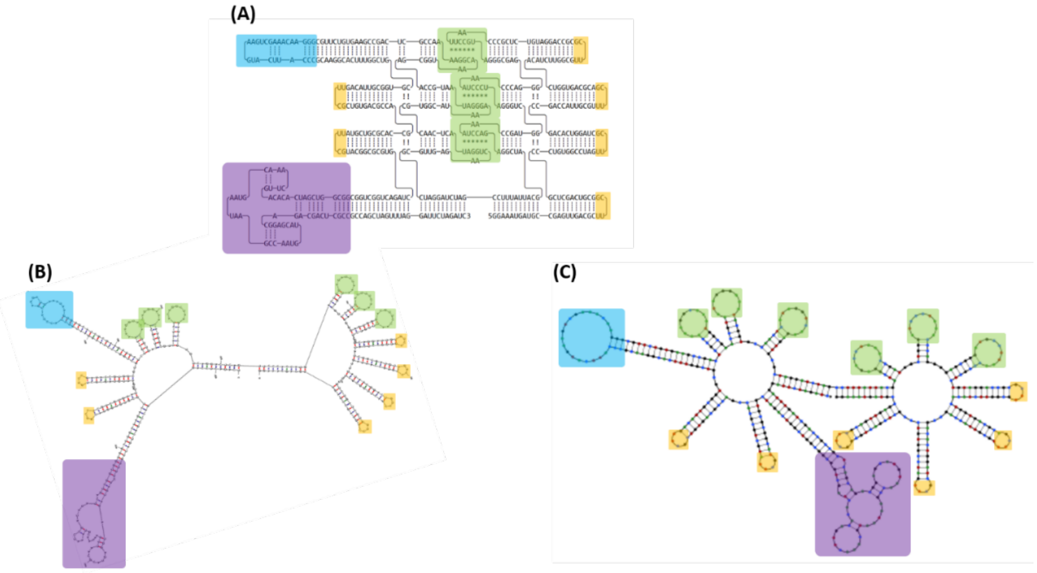


Figure S6 (A) 2D illustration of 4H-2NN1 RNA origami anticoagulant. (B and C) Computational analysis of RNA origami folding analyzed by Mfold RNA and NUPACK, online software. Blue and purple rectangles represent exosite-I, and exosite-II-binding aptamers, respectively. Yellow and green rectangles indicate tetra loops and kissing loop motifs.

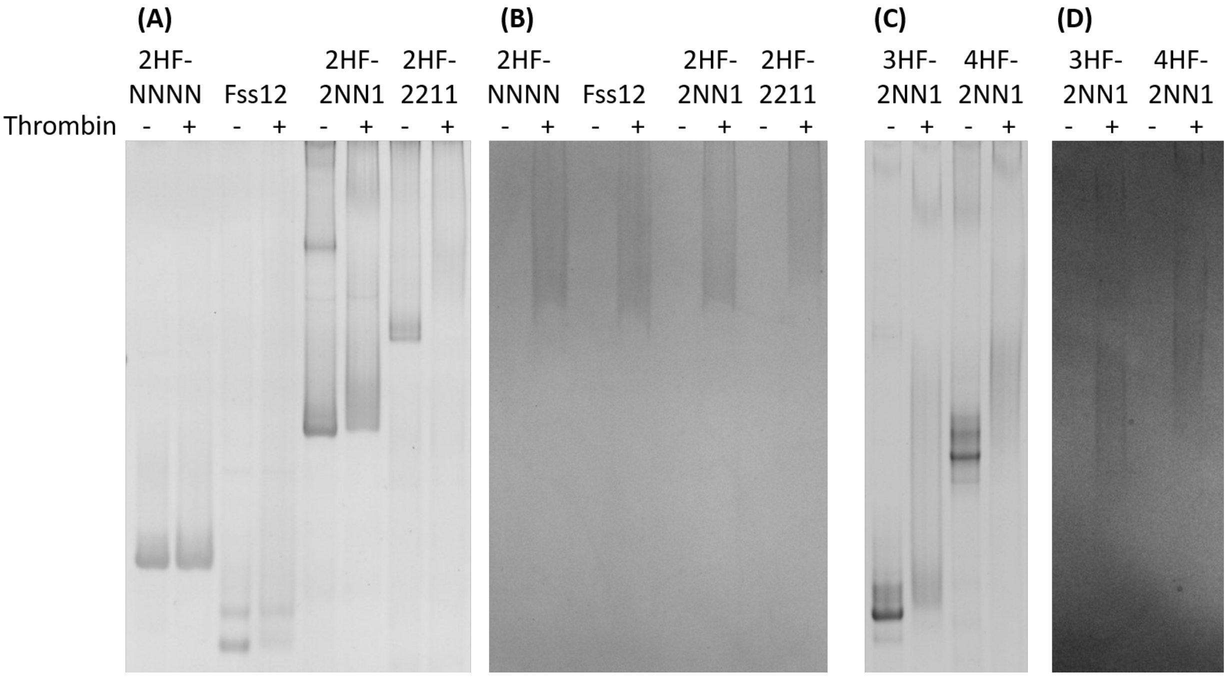


**Figure S7** Binding of RNA origami with thrombin by 6% native acrylamide gel electrophoresis. (A) RNA origami incubated with thrombin at 37 C for 1 hr before characterization. Gel A and B are the same gel and run at 150 V for 3 hr. Gel C and D are the same gel and run at 150 V for 6 hr. (A and C) Nucleic acid-stained gel, ethidium bromide. (B and D) Protein-stained gel, Coomassie blue. Negative and positive values indicated absence and presence of thrombin, respectively.

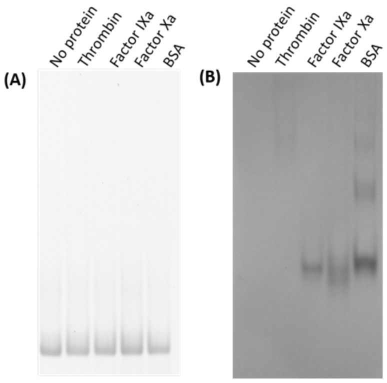


**Figure S8** Specific binding test of 2HF-NNNN (no aptamers) with four different proteins by native acrylamide gel electrophoresis. Lane 1: 2HF-NNNN. Lane 2: 2HF-NNNN incubated with thrombin. Lane 3: 2HF-NNNN incubated Factor IXa. Lane 4: 2HF-NNNN incubated Factor Xa, and Lane 5: 2HF-NNNN incubated BSA. 6% native PAGE gel was run at 150 for 3 hr. (A) Nucleic-acid stained gel, ethidium bromide. (B) Protein-stained gel, coomassie blue. (A and B) Both are the same gel

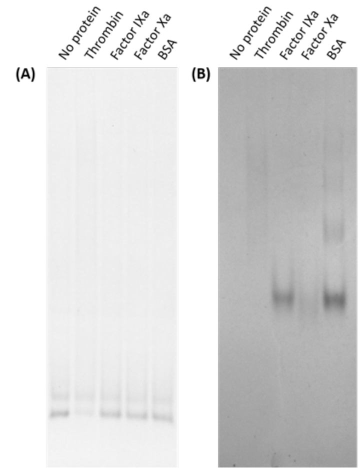


**Figure S9** Specific binding test of 31nt-linked two aptamers (Fss12) with four different proteins by native acrylamide gel electrophoresis. Lane 1: Fss12. Lane 2: Fss12 incubated with thrombin. Lane 3: Fss12 incubated Factor IXa. Lane 4: Fss12 incubated Factor Xa, and Lane 5: Fss12incubated BSA. 6% native PAGE gel was run at 150 for 3 hr. (A) Nucleic-acid stained gel, ethidium bromide. (B) Protein-stained gel, coomassie blue. (A and B) Both are the same gel

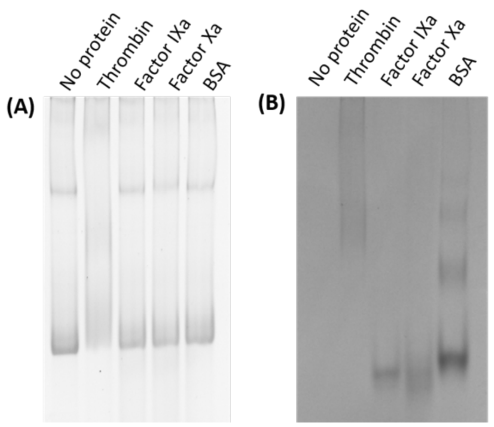


**Figure S10** Specific binding test of 2HF-2NN1 with four different proteins by native acrylamide gel electrophoresis. Lane 1: 2HF-2NN1. Lane 2: 2HF-2NN1 incubated with thrombin. Lane 3: 2HF-2NN1 incubated Factor IXa. Lane 4: 2HF-2NN1 incubated Factor Xa, and Lane 5: 2HF-2NN1 incubated BSA. 6% native PAGE gel was run at 150 for 3 hr. (A) Nucleic-acid stained gel, ethidium bromide. (B) Protein-stained gel, coomassie blue. (A and B) Both are the same gel

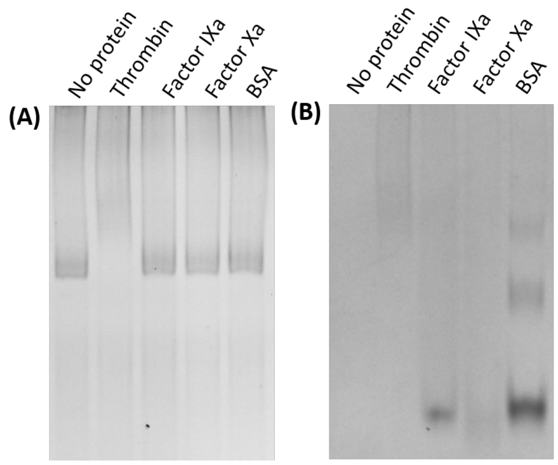


**Figure S11** Specific binding test of 2HF-2211 with four different proteins by native acrylamide gel electrophoresis. Lane 1: 2HF-2211. Lane 2: 2HF-2211 incubated with thrombin. Lane 3: 2HF-2NN1 incubated Factor IXa. Lane 4: 2HF-2211 incubated Factor Xa, and Lane 5: 2HF-2211 incubated BSA. 6% native PAGE gel was run at 150 for 3 hr. (A) Nucleic-acid stained gel, ethidium bromide. (B) Protein-stained gel, coomassie blue. (A and B) Both are the same gel

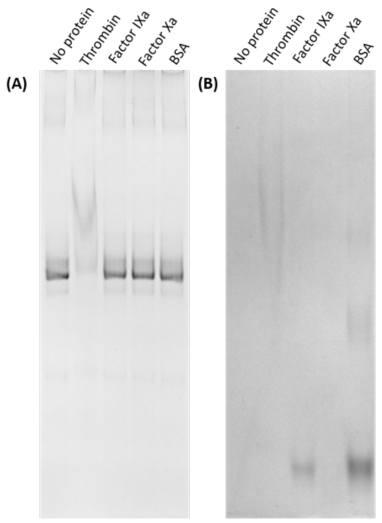


**Figure S12** Specific binding test of 3HF-2NN1 with four different proteins by native acrylamide gel electrophoresis. Lane 1: 3HF-2NN1. Lane 2: 3HF-2NN1 incubated with thrombin. Lane 3: 3HF-2NN1 incubated Factor IXa. Lane 4: 3HF-2NN1 incubated Factor Xa, and Lane 5: 3HF-2NN1 incubated BSA. 6% native PAGE gel was run at 150 for 6 hr. (A) Nucleic-acid stained gel, ethidium bromide. (B) Protein-stained gel, coomassie blue. (A and B) Both are the same gel.

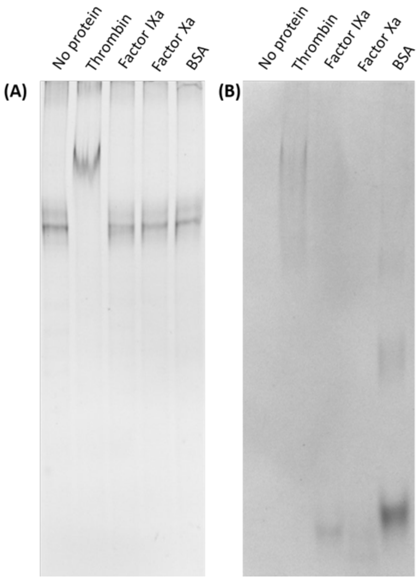


**Figure S13** Specific binding test of 4HF-2NN1 with four different proteins by native acrylamide gel electrophoresis. Lane 1: 2HF-4NN1. Lane 2: 4HF-2NN1 incubated with thrombin. Lane 3: 4HF-2NN1 incubated Factor IXa. Lane 4: 2HF-4NN1 incubated Factor Xa, and Lane 5: 4HF-2NN1 incubated BSA. 6% native PAGE gel was run at 150 for 6 hr. (A) Nucleic-acid stained gel, ethidium bromide. (B) Protein-stained gel, coomassie blue. (A and B) Both are the same gel

**Pyrimidine modification levels**

The production of 2’-fluoro-modified RNA origami is expensive due to the higher price of unnatural NTPs, 2’-F dUTP and 2’-F dCTP. In an attempt to reduce the production costs, we transcribed RNA origami with various ratios of native nucleotide (CTP and UTP) and modified nucleotides (2’-F dCTP and 2’-F dUTP). The transcribed RNA origami was characterized by denaturing PAGE. As a result, we successfully produced fully-modified, partially-modified (75%, 50% and 25%) and non-modified RNA origami. Unfortunately, all levels of partially- and non-modified RNA origami were shown to be unstable in RNase A (see Fig S14). However, we showed that the fully-modified (all pyrimidines modified) RNA is stable in the presence of nucleases. Therefore, we continued further experiments with all-pyrimidine-modified RNA origami.

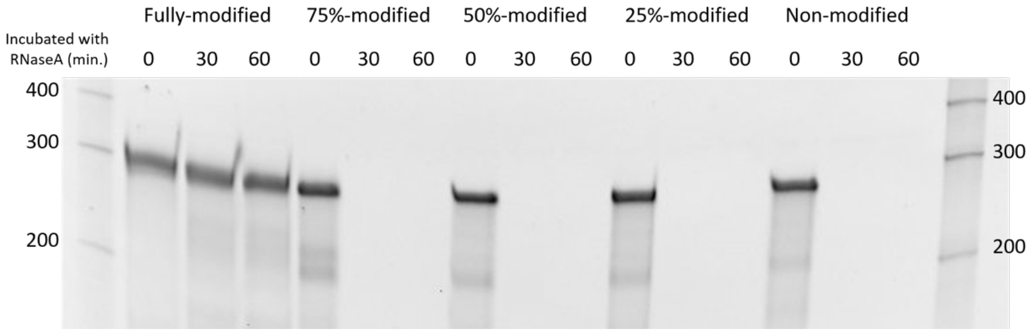


**Figure S14** Stability test of RNA origami against RNase A (10 μg/ml). RNA origami incubated with RNase A at 37 C for 30 and 60 minutes. For the control samples, RNA origamis were incubated at 37 C without RNase A. Stability of RNA origami after treated with RNase A was characterized by denaturing PAGE gel and post-stained with ethidium bromide. Fully-modified RNA origami contains 2’F-dCTP, 2’F-dUTP, ATP, and GTP at ratio of 1:1:1:1. For non-modified RNA origami, the 2’fluoro-dCTP and 2’fluoro-dUTP were substituted with CTP and UTP. In case of 75%-modified RNA origami, we mixed the 2’fluoro-dCTP with CTP and 2’fluoro-dCTP with CTP at ratio of 3:1. Lane 1 and 17 are ssRNA marker.

**Table S1** DNA template and DNA primer sequences

| Name | Sequence (5’ ----> 3’) |
|---|---|
| 2H-Gblock-NNNN | CACTTTCAGCCCTCTTATCCTCGGCGGATCCTTCTAATACGAC<br>TCACTATAGGGAGATCGAGCGACTTCCGACTTCGGTCGGGAG<br>TCGGGCTAGTCATCTTCGGATGATTAGCCGCTGGTGAAGCCT<br>CCACGCCAGCCTCGGTCTCCCGCAGTAGGATCGGACTGAAGG<br>AGGCACGGTCCCAGCCGAAGTGTCTTGCTTCGGCAAGGCACT<br>TTGGCTGCTAGACTGGCTGGCTTCGGCCAGCTAGTTTAGGATT<br>CTATTGC |
| 2H-Gblock-2NN1 | CACTTTCAGCCCTCTTATCCTCGGCGGATCCTTCTAATACGAC<br>TCACTATAGGGAGATCGAGCGACTTCCGACTTCGGTCGGGAG<br>TCGGGCTAGTCATCTTCGGATGATTAGCCGCTGGTGAAGCCT<br>CCACGCCAGCCTCGGTCTCCCGCAGTAGGATCGGACTGAAGG<br>AGGCACGGTCCCAGCCGAAGTGTCTTGCGGGAACAAAGCTGA<br>AGTACTTACCCGCAAGGCACTTTGGCTGCTAGACTGGCTGGC<br>GGCGGTCGATCACACAGTTCAAACGTAATAAGCCAATGTACG<br>AGGCAGACGACTCGCCGCCAGCTAGTTTAGGATTCTATTGC |
| 31nt-linked-aptamers-Gblock | CACTTTCAGCCCTCTTATCCTCGGCGGATCCTTCTAATACGAC<br>TCACTATAGGGAACAAAGCTGAAGTACTTACCCACCTTACCA<br>CTCCACCTCACTCACCTATTACGGCGGTCGATCACACAGTTCA<br>AACGTAATAAGCCAATGTACGAGGCAGACGACTCGCC |
| 3H-Gblock-2NN1 | CGGCCAGTGAATTCGAGCTCGGTACCCGGGAGATCTCACTTT<br>CAGCCCTCTTATCCTCGGCCGATCCTTCTAATACGACTCACTA<br>TAGGAAATGATGCCGAGTTGACGCTTCGGCGTCAGCTCGCCC<br>TGTGGCCTAGTTCGCTAGGTCACAGACATCTTGGCGTTCGCGC<br>CAGGATGTCTCGCCCAATTCCGTAGGGCGAGGGTAGCCAAAT<br>CCAGAGGCTAGCATTATTTCCGATCTAGGATCGCGTTGAGAA<br>CTGGATACTCAACAGCGGTAAACGGAAAACCGCTCAGCCGA<br>AGTGTCTTGCGGGAACAAAGCTGAAGTACTTACCCGCAAGGC<br>ACTTTGGCTGGCCACGCGTCGTATTCGTACGGCGCGTGCTAG<br>ACTGGCTGGCGGCGGTCGATCACACAGTTCAAACGTAATAAG<br>CCAATGTACGAGGCAGACGACTCGCCGCCAGCTAGTTTAGGA<br>TTCTAGATCTCTCTAGAGTCGACCTGCAGGCATGCAA |

| Name | Sequence (5’ ----> 3’) |
|---|---|
| 4H-Gblock-2NN1 | CGGCCAGTGAATTCGAGCTCGGTACCCGGGAGATCTCACTTT<br>CAGCCCTCTTATCCTCGGCCGATCCTTCTAATACGACTCACTA<br>TAGGAAATGATGCCGAGTTGACGCTTCGGCGTCAGCTCGCCC<br>TGTGGCCTAGTTCGCTAGGTCACAGCCGACCATTGCGTTTCGA<br>CGCAGTGGTCACATCTTGGCGTTCGCGCCAGGATGTCTCGCC<br>CAATTCCGTAGGGCGAGGGGACCCAAATCCCTAGGGTCGGTA<br>GCCAAATCCAGAGGCTAGCATTATTTCCGATCTAGGATCGCG<br>TTGAGAACTGGATACTCAACCGTGGCATAAAGGGATAATGCC<br>AAGCGGTAAACGGAAAACCGCTCAGCCGAAGTGTCTTGCGG<br>GAACAAAGCTGAAGTACTTACCCGCAAGGCACTTTGGCTGCG<br>TGGCGTTACAGTTCGCTGTGACGCCAGCCACGCGTCGTATTC<br>GTACGGCGCGTGCTAGACTGGCTGGCGGCGGTCGATCACACA<br>GTTCAAACGTAATAAGCCAATGTACGAGGCAGACGACTCGCC<br>GCCAGCTAGTTTAGGATTCTAGATCTCTCTAGAGTCGACCTGC<br>AGGCATGCAAGCT |
| 2H-Gblock-2211_Fragment1 | CACTTTCAGCCCTCTTATCCTCGGCGGATCCTTCTAATACGAC<br>TCACTATAGGGAGATCGAGCGACTTCCGACTCTGGCGGTCGA<br>TCACACAGTTCAAACGTAATAAGCCAATGTACGAGGCAGACG<br>ACTCGCCAGAGTCGGGAGTCGGGCTAGTCATCAGGCACGGGA<br>ACAAAGCTGAAGTACTTACCCGTGCCTGATGATTAGCCGCTG<br>GTGAAGCCTCCAC |
| 2H-Gblock-2211_Fragment2 | CTGATGATTAGCCGCTGGTGAAGCCTCCACGCCAGCCTCGGT<br>CTCCCGCAGTAGGATCGGACTGAAGGAGGCACGGTCCCAGCC<br>GAAGTGTCTTGCGGGAACAAAGCTGAAGTACTTACCCGCAAG<br>GCACTTTGGCTGCTAGACTGGCTGGCGGCGGTCGATCACACA<br>GTTCAAACGTAATAAGCCAATGTACGAGGCAGACGACTCGCC<br>GCCAGCTAGTTTAGGATTCTATTGC |
| Exosite-I aptamer DNA template | TCGGCGGATCCTTCTAATACGACTCACTATAGGCGGTCGATC<br>ACACAGTTCAAACGTAATAAGCCAATGTACGAGGCAGACGA<br>CTCGCC |
| Exosite-II aptamer DNA template | TCGGCGGATCCTTCTAATACGACTCACTATAGGGAACAAAGC<br>TGAAGTACTTACCC |
| 2H_Forward Primer | CACTTTCAGCCCTCTTATCC |
| 2H_Reverse Primer | GCAATAGAATCCTAAACTAGCTGG |
| 31nt-apt_Forward primer | CACTTTCAGCCCTCTTATCC |
| 31nt-apt_Reverse primer | GGCGAGTCGTCTG CC |
| 3 and 4H_Forward Primer | AGATCTCACTTTCAGCCCTCTTATC |
| 3 and 4H_Reverse Primer | GATCTAGAATCCTAAACTAGCTGG |
| Exosite-1 antidote | GTCTGCCTCGTACATTGGCT |
| Exosite-2 antidote | GTACTTCAGCTTTGTTCCC |
